# Identification of a Conserved B-Cell Epitope on the Capsid Protein of Porcine Circovirus Type 4

**DOI:** 10.1101/2024.03.18.585584

**Authors:** Zheng Fang, Mingxia Sun, Xuehui Cai, Tongqing An, Yabin Tu, Haiwei Wang

## Abstract

Porcine circovirus type 4 (PCV4), a recently identified circovirus, is prevalent in numerous provinces in China, as well as in South Korea, Thailand, and Europe. PCV4 virus rescued from an infectious clone showed pathogenicity, suggesting the economic impact of PCV4. However, there remains a lack of understanding regarding the immunogenicity and epitopes of PCV4. This study generated a monoclonal antibody (MAb) 1D8 by immunizing mice with PCV4 virus-like particles (VLPs). Subsequently, the epitope recognized by the MAb 1D8 was identified by truncated protein expression and alanine scanning mutagenesis analysis. Results showed that the ^225^PKQG^228^ located at the C-terminus of the PCV4 Cap protein is the minimal motif binding to the MAb. Homology modeling analysis and immunoelectron microscopy revealed that the epitope extends beyond the outer surface of the PCV4 VLP. Moreover, the epitope is highly conserved among PCV4 strains and does not react with other PCVs. Together, the MAb 1D8 recognized epitope shows potential for detecting PCV4. These findings significantly contribute to the design of antigens for PCV4 detection and control strategies.

**IMPORTANCE:** Porcine circovirus type 4 (PCV4) is a novel circovirus. Although PCV4 has been identified in several countries, including China, Korea, Thailand, and Spain, no vaccine is available. Given the potential pathogenic effects of PCV4 on pigs, PCV4 could threaten the global pig farming industry, highlighting the urgency for further investigation. Thus, epitopes of PCV4 remain to be determined. Our finding of a conserved epitope significantly advances vaccine development and pathogen detection.

## INTRODUCTION

Porcine circovirus (PCV) is a member of the genus Circovirus within the family Circoviridae. Since the first discovery of the circular single-stranded DNA virus in pigs in 1974, four PCVs have been progressively reported[1, 2, 3, 4, 5, 6]. Porcine circovirus type 2 (PCV2) is firmly associated with PCV-associated diseases (PCVADs)[7, 8, 9], causing significant economic losses in pig farming. Despite the wide distribution of PCV3, its pathogenicity remains controversial[10]. Porcine circovirus type 4 (PCV4) was first identified in Hunan, China, in 2019[6]. It has typical PCV characteristics, such as a conserved nine-nucleotide hairpin structure and two major open reading frames (ORFs)[11, 12]. Given the historical impact of PCVs, further studies are imperative to determine whether PCV4 presents a potential threat to the pig industry. Previous reports indicated the widespread presence of PCV4 across numerous provinces in China, with detections in South Korea and Thailand[6, 12, 13, 14, 15, 16, 17, 18, 19, 20]. In addition, PCV4 was recently detected in European wild boars[21]. Notably, PCV4 rescued from an infectious clone showed pathogenicity in pigs[22]. Considering the historical impact of PCVs and their widespread distribution across various regions, further epidemiological research is necessary to clarify the potential threat of PCV4 to the pig industry.

The PCV4 virion is assembled by Cap protein and viral genome. PCV Cap protein is the only structural protein with multiple functions. Extensive research on the PCV2 Cap protein has been conducted, including its immunogenicity and its interaction with the host[23, 24, 25, 26, 27, 28]. PCV4 Cap protein has been reported to interact with DDX21 and localize to the nucleus, a characteristic shared with other PCV Caps[29]. Additionally, PCV4 VLP has a comparable structure and morphology to other PCVs, measuring 20 nm[30, 31]. PCV4 VLP was also shown to enter PK-15 and 3D4/21 Cells, consistent with permissive cell lines of PCV2[22, 30, 32, 33]. Nevertheless, PCV4 has less than 50% identity of amino acid sequences compared to other PCVs, exhibiting significant differences in sequence and specific structure[6, 30]. The immunogenicity and the dominant epitope of PCV4 remain to be investigated. VLP has been widely used as a tool for vaccine design and immunogenicity studies due to its structural and immunological similarities to original viruses[34, 35]. For instance, PCV2 VLP was used to evaluate its immunogenicity and identify the epitopes. PCV2 epitopes recognized by MAb 3H11 were located on PCV2 VLP[25]. Therefore, VLP can also serve as a valuable tool for studying the epitopes and immunogenicity of PCV4.

In this study, we first generated and characterized seven monoclonal antibodies specifically targeting PCV4 Cap, which are crucial for type-specific detecting PCV4. Furthermore, we finely mapped the epitope to the C-terminus recognized by the MAb 1D8 by GST-fusion protein expression and Western blot analysis. Moreover, this epitope is found to be conserved in PCV4 but not found in other pig-derived pathogens, which supports the use of the MAb 1D8 for the type-specific detection of PCV4. Based on corresponding studies on PCV2, we also discussed the potential significance of this unique epitope. Our findings could be used to design antigens for PCV detection and prevention.

## MATERIALS AND METHODS

### Cells, plasmids

The PK-15 and SP2/0 myeloma cells were cultured in Dulbecco’s modified Eagle’s medium (DMEM, Gibco, USA) supplemented with 10% fetal bovine serum (FBS, ExCell, Australia). They were all cultured in a humidified incubator at 37°C and 5% CO_2_ atmosphere. The PCI-tb201, pET28a, and pGEX-6p-1 vectors were preserved in our laboratory before.

### Expression and purification of PCV4 VLPs

The PCV4 VLPs were prepared in our laboratory before[31]. PCV4 Cap gene based on the sequence from Genbank (Accession No. MT769268.1) was codon-optimized and synthesized by Ruibiotech (Beijing). Then, it was ligated into the vector pET28a. The recombinant plasmid was transfected into *E. coli* BL21 (DE3) cells (Vazyme, China) to express PCV4 Cap. Assembled PCV4 VLPs were purified through Rose Plus Q XP column (Nanomicrotech, China), followed by Sepharose 6FF 16/96 column (Bestchrom, China), which were connected to a FPLC system (ÄKTA avant, Cytiva, Sweden). The PCV2-VLPs[36] and PCV3-VLPs[37] were previously prepared in our lab.

### Preparation of Monoclonal Antibodies

Six-week-old female BALB/c mice were purchased from the Experimental Animal Center of Harbin Veterinary Research Institute (Chinese Academy of Agricultural Sciences, China). They were immunized with 20 µg of PCV4 VLPs with an equal volume of adjuvant Montanide ISA-201 (SEPPIC, France). After the initial immunization, two boosters with the same PCV4 VLPs and adjuvant doses were given at 14-day intervals. Following three times vaccinations, antibody titers against PCV4 VLPs in serum reached 1:51200 as measured by PCV4-VLP-ELISA[31]. One week after the last immunization, the mice were intraperitoneally boosted with 20 µg of PCV4 VLPs without adjuvant. The spleen cells were separated and fused with SP2/0 myeloma cells three days later by 50% (w/v) PEG4000 (Sigma-Aldrich, USA). PCV4-VLP-ELISA screened the antibodies in supernatants of hybridoma. Then, the reactivity with PCV4 Cap was verified by indirect immunofluorescence assay (IFA) and Western blot. Positive hybridoma were subcloned four times. Antibody subtype identification was performed using the SBA ClonotypingTM System/HRP Kit (Southern Biotech, Birmingham, AL, USA). To minimize the background impurities observed during immunoelectron microscopy, we purified MAb 1D8. Mice were selected and injected with liquid paraffin to induce an immune response in the peritoneal cavity. Monoclonal hybridoma cells 1D8 were cultured. After one week, hybridoma cells were injected into the same cavity to yield a substantial quantity of monoclonal antibodies. The ascites were then centrifuged, and the supernatant was subjected to HiTrap Protein G HP (GE, USA) to obtain purified MAb 1D8.

### SDS-PAGE and Western blot

Approximately equal amounts of GST fusion proteins and alanine-mutated GST fusion proteins were separated by 12% SDS-PAGE and transferred to PVDF transfer membranes (Millipore, MA, USA). The membranes were blocked with 5% skimmed milk powder for 2 hours at 37°C. Hybridoma cell supernatant was used as the primary antibody and incubated for 1 hour at 37°C, followed by three washes with PBST. Then HRP-conjugated goat anti-mouse IgG (Sigma, diluted 1:10,000, USA) was added. The membranes were incubated for 1 hour at 37°C before being scanned using a near-infrared fluorescence scanning imaging system (Odyssey CLX, USA).

### Cloning and expression of the full-length and truncated PCV4 Cap proteins

To identify the epitope recognized by MAb 1D8, the full-length Cap proteins and overlapping peptides spanning the Cap protein were expressed. The full-length PCV4 Cap protein and overlapping peptides sequence was amplified from the PCV4 VLP expression plasmid. For the PCR reaction, 25 µL mixture was prepared containing 12.5 µL of 2× PCR Buffer for KOD FX, 5 µL of 2 mM deoxyribonucleotide triphosphate (dNTP), 0.75 µL of forward primer, 0.75 µL of reverse primer, 0.5 µL of KOD FX (TOYOBO CO., LTD, Japan), and 5 µL of sterile ddH2O. The PCR reaction was subjected to pre-denaturation at 94°C for 4 minutes, followed by 30 cycles of denaturation at 98°C for 30 seconds, annealing at 56°C for 30 seconds, and extension at 68°C for 3 minutes. A final extension was performed at 68°C for 10 minutes. After determining that the reactivity was located in the GST-Cap214-228 fragment, the fragment was truncated for expression one by one from each terminus respectively. Complementary primer pairs containing EcoRI and XhoI restriction sites in each GST-fused fragment’s oligonucleotide coding were synthesized and annealed. All the primer information used for cloning each fragment were displayed (Table 1-2). Then, the annealed primers were inserted into the pGEX-6p1 vector. SDS-PAGE and Western blot were used to analyze the GST-fusion proteins.

**TABLE 1.**
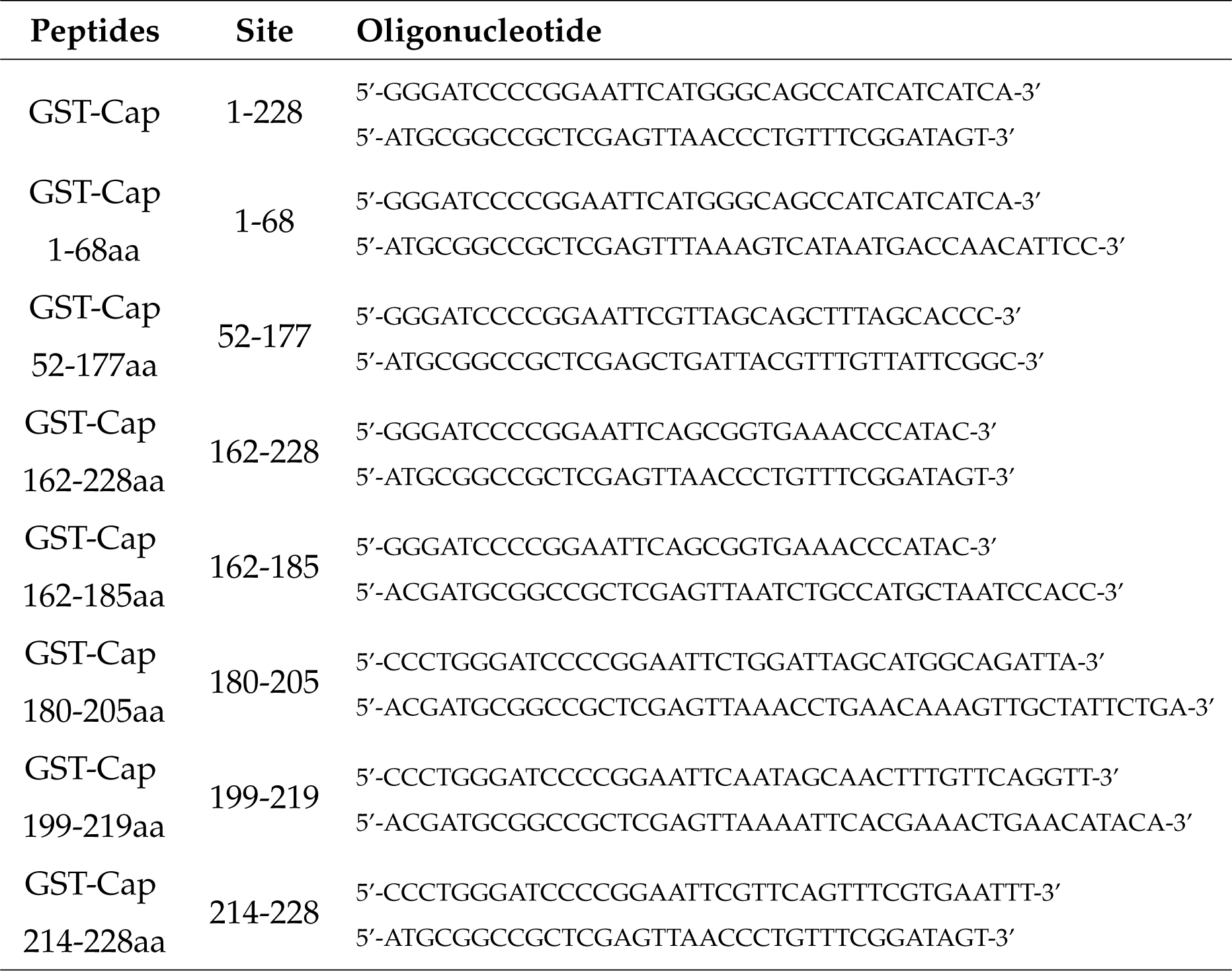
Complementary oligonucleotides coding for the truncated GST-Cap segments.

**TABLE 2.**
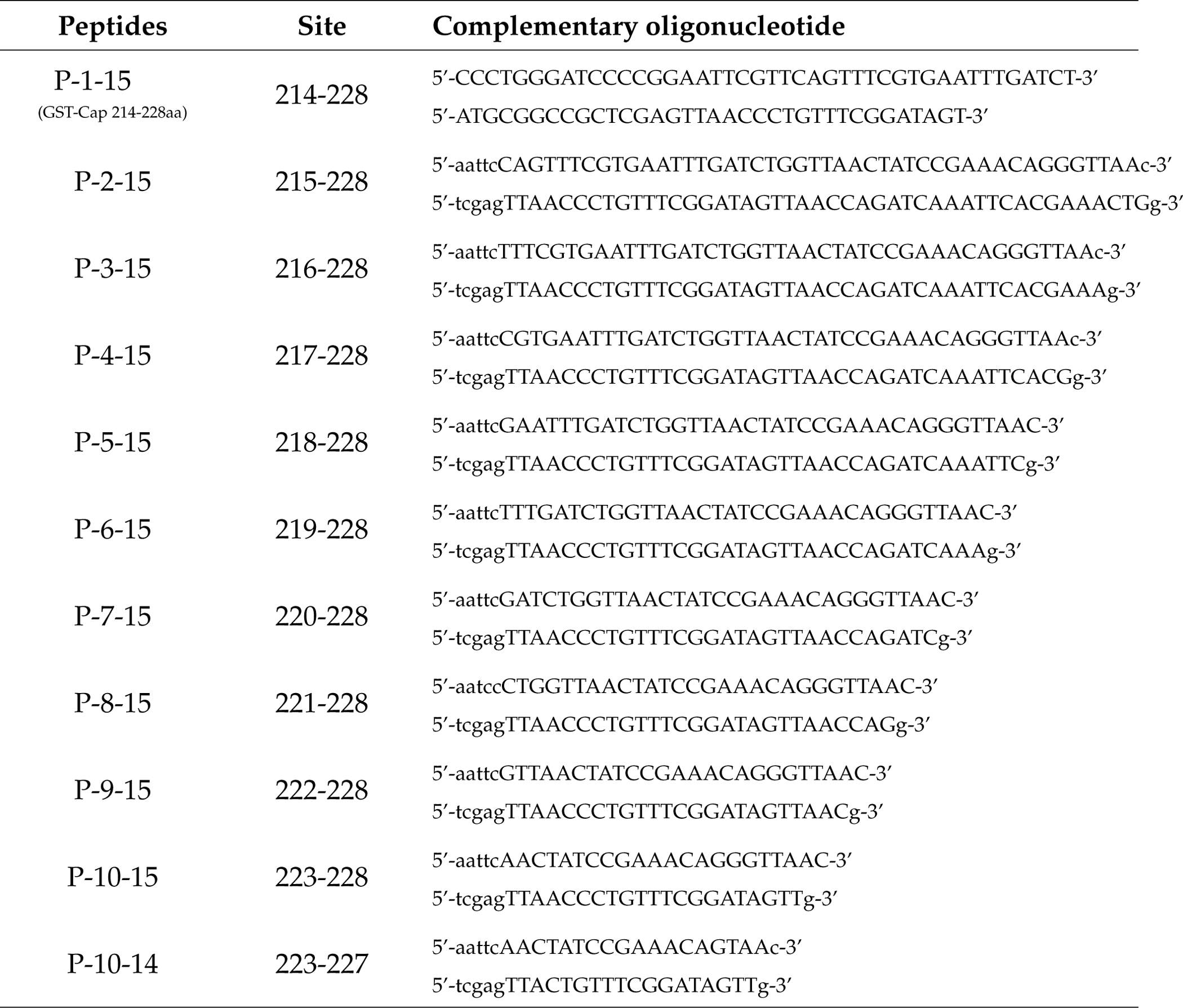
Complementary oligonucleotides coding for the truncated GST-Cap214-228aa segments.

### Indirect immunofluorescence assay

The PCV4 Cap protein sequence was amplified and inserted into the PCI-tb201 vector. Following transfection of PK-15 cells with 200 ng of the recombinant plasmid using jetPRIME transfection reagent (Polyplus, USA), the cells were cultured for 48 hours. After rinsing with PBS, the cells were fixed in 4% paraformaldehyde for 20 minutes, treated with 0.2% Triton X-100 for 30 minutes at room temperature, and blocked with 1% BSA for 30 minutes. Hybridoma supernatant (200 µL) was added and incubated at 37°C for 1 hour. After rinsing with PBS, we added fluorescein isothiocyanate (FITC)-conjugated goat anti-mouse IgG (Sigma, USA) at a dilution of 1:200 and incubated it at 37°C for 1 hour. The plates were washed with PBS and observed under a fluorescence microscope (EVOS F1, AMG, USA).

### Alanine-scanning mutagenesis

To identify the crucial residues in the epitope necessary for the MAb 1D8 binding, we created nine alanine mutants that targeted the epitope ^220^DLVNYPKQG^228^. Paired primers were used to introduce these mutations (Table 3), which were then synthesized and annealed into the pGEX-6P-1 vector for expression in E.coli. After expression, bacterial lysate supernatants were collected and analyzed using SDS-PAGE and Western blot.

**TABLE 3.**
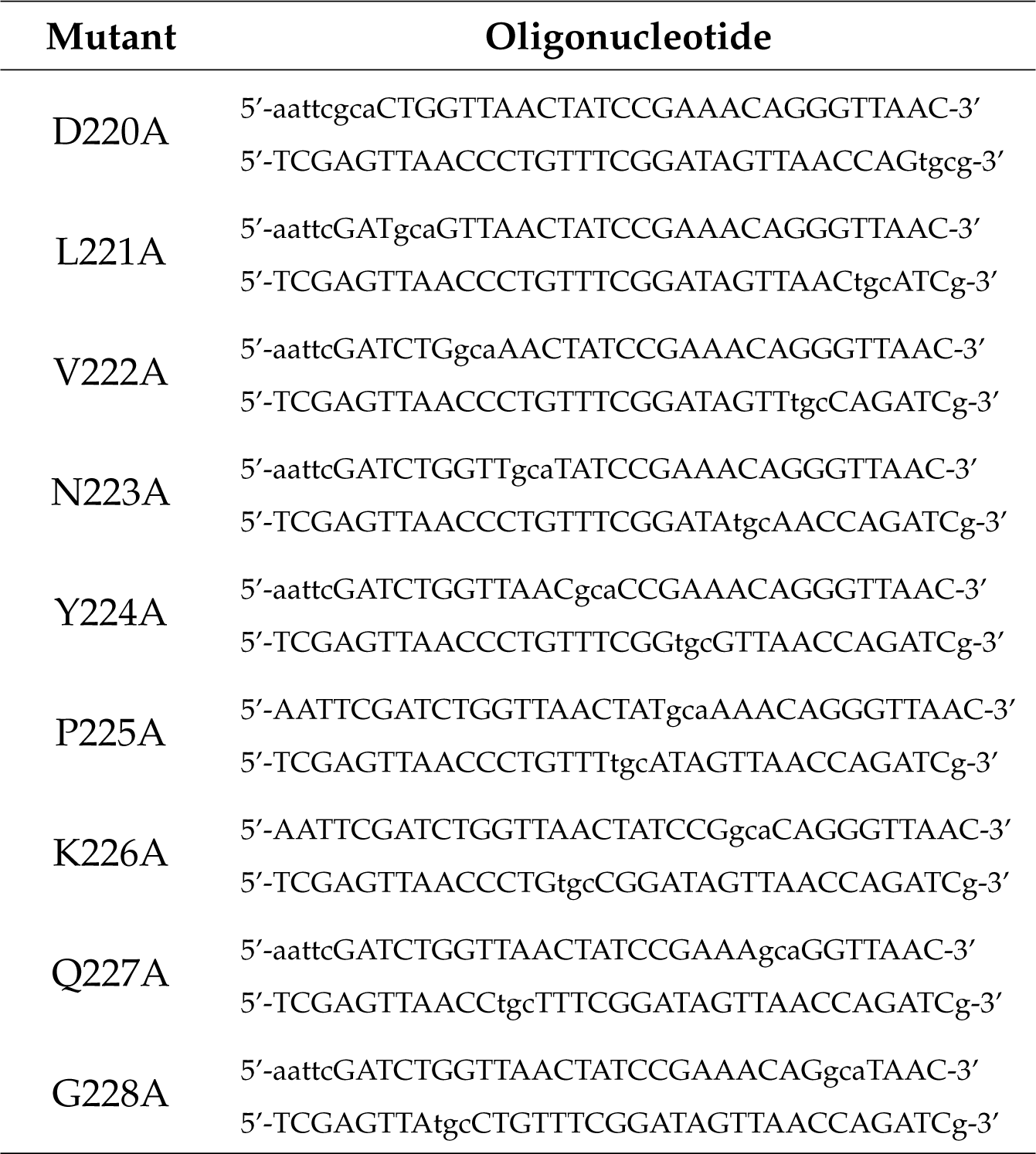
Primers used to generate specific mutations in the Cap gene.

### Immunoelectron microscopy

Transmission electron microscopy (TEM) was used to visualize the binding complex between PCV4 VLPs and MAb 1D8. PCV4 VLP samples were initially deposited onto copper grids for 3 minutes. They were then incubated with 3% BSA for 30 minutes, followed by immersion in purified MAb 1D8 and incubation for 40 minutes. Rinse the copper mesh with distilled water for three minutes, and then rinse it twice. A 5 nm colloidal gold-conjugated antibody (diluted 1:100, Sigma, USA) was applied to the grid. After 40 minutes of incubation at room temperature, the grid was rinsed three times with distilled water for 3 minutes each time. Negative staining was performed by incubating the samples in 2% phosphotungstic acid for 20 seconds. After drying, the samples were observed using a TEM instrument (Hitachi, Tokyo, Japan).

### Biological information analysis

The amino acid sequence of the PCV4 Cap protein was obtained from the Uniprot database (ID: A0A7M4CFD7). The secondary structure of the Cap was predicted using the online secondary structure analysis software PSIPRED (http://bioinf.cs.ucl.ac.uk/psipred/). Subsequently, epitopes of the PCV4 Cap protein were predicted using Bepipred (https://services.healthtech.dtu.dk/services/BepiPred-3.0/).

Templates of the Cap protein and VLP were selected, and homology modeling of VLP was performed using the SWISS-MODEL (https://swissmodel.expasy.org/interactive) online software. Homology modeling of Cap was performed using the I-TASSER server (https://zhanggroup.org/I-TASSER/). The 3D structures of the Cap protein and VLP were then visualized and analyzed using Chimera X.

We compared the representative sequences to analyze the conservation of ^225^PKQG^228^ epitope in PCVs. PCVs Cap sequences were retrieved from the NCBI GenBank database (http://www.ncbi.nlm.nih.gov/). Representative strains were selected for each genotype, and amino acid sequences were compared using the ESPript 3.0 online tool (http://espript.ibcp.fr/ESPript/ESPript/).

## RESULTS

### Insights of Cap Protein from Bioinformatics Analysis

To study the B-cell epitopes precisely, we predicted the secondary structure of the protein using the online software PSIPRED (http://bioinf.cs.ucl.ac.uk/psipred/). The PCV4 Cap protein is composed of 228 amino acid residues. The Cap protein is composed of 3.90% alpha-helix region, 33.33% beta-sheet region, and 62.70% randomized coils (Fig.1A). The prediction of epitopes for the Cap was conducted using the BepiPred3.0 tool (https://services.healthtech.dtu.dk/services/BepiPred-3.0/). A threshold score of 0.2 was applied to the BepiPred epitopes to identify the potential B-cell epitopes (Fig.1B). The result showed that the PCV4 Cap has eight potential B-cell epitopes (Table 4). Most of the epitopes were located in the coil structure, with a gravy lower than 0, indicating better hydrophilicity.

**FIG 1.**
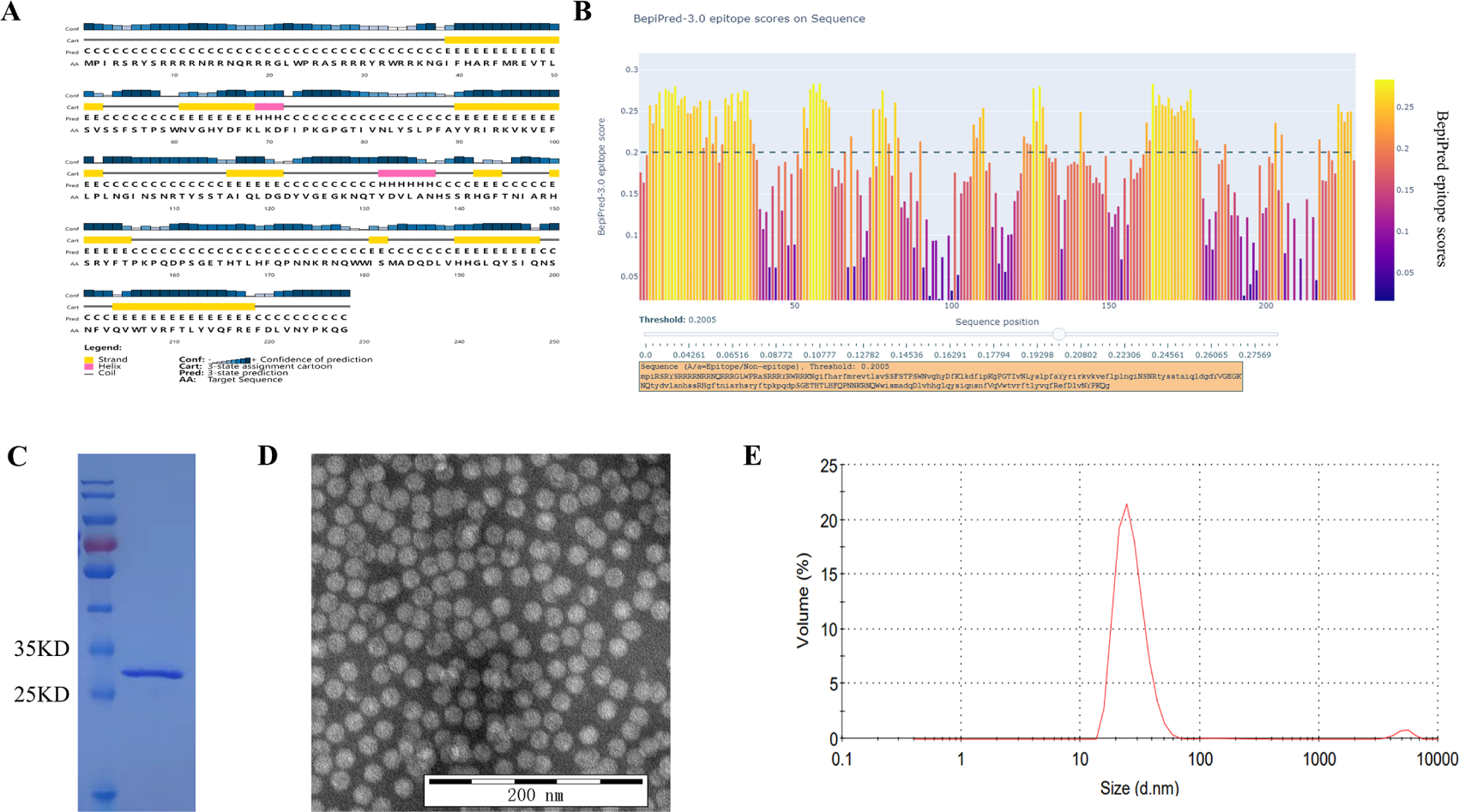
Characterization of PCV4 Cap Protein and immunized VLP. (A) Secondary structure prediction analysis of PCV4 Cap protein using PSIPRED online software. (B) Prediction of epitopes for PCV4 Cap B-cell using BepiPred3.0 tool. A threshold score of 0.2 was applied to identify potential B-cell epitopes. (C) SDS-PAGE analysis of purified PCV4 VLPs, which were intended for immunization. (D) TEM image showing the morphology of immunized PCV4 VLPs. (E) DLS analysis results display the size distribution of immunized PCV4 VLPs, indicating an average diameter of approximately 20nm.

**TABLE 4.** Predicted B-cell epitopes on the Cap protein.

| Epitope | Amino acid sequence | Position(aa) | Secondary structure |
| --- | --- | --- | --- |
| B-1 | RSRYSRRRRNRRNQRRRGWPR | 4-25 | 100% coil |
| B-2 | SRRRYRWRRKN | 27-37 | 100% coil |
| B-3 | SSFSTPSWN | 53-61 | 89% coil and 11% strand |
| B-4 | KGPGTIVNL | 75-83 | 100% coil |
| B-5 | NSNR | 107-110 | 100% coil |
| B-6 | YVGEGKNQ | 123-130 | 100% coil |
| B-7 | SGETHTLHFQPNNKRNQW | 162-179 | 100% coil |
| B-8 | NYPKQ | 223-227 | 100% coil |

### Expression and purification of PCV4 VLP

The PCV4 Cap proteins were successfully expressed using the prokaryotic system. The supernatant was collected and purified using two-step chromatography[31]. The purified protein exhibited a molecular weight of approximately 27 kDa, as shown by the SDS-PAGE (Fig. 1C). TEM and DLS analyses revealed a significant quantity of fully assembled virus-like particles, which were approximately 20 nm in size (Fig. 1D-E). Finally, we successfully produced pure assembled PCV4 VLPs.

### Preparation of MAbs against PCV4 VLPs

Mice were immunized three times with PCV4 VLPs, and the antibodies in their serum against PCV4 Cap were quantified positive at a dilution of 1:51200. Following immunization, splenocytes were isolated and fused with SP2/0 cells. Subsequently, hybridoma cell supernatants were assessed using PCV4-VLPs-ELISA. Seven hybridoma clones (1D6, 1D8, 3F10, 5E10, 3F11, 2F7, and 5D10) were found as producers of MAbs against the PCV4 Cap protein. These cloned cells showed stability in antibody secretion through four subclones.

Seven MAbs were identified to react specifically with PCV4 Cap protein through IFA and Western blot (Fig.2). Subtyping analysis revealed a consistent *κ* light chain and IgG2a heavy chain among all seven MAbs. The subcloned hybridoma cells were injected into the peritoneal cavity of mice, and monoclonal antibodies were obtained by purifying ascites using a Protein G affinity chromatography column.

**FIG 2.**
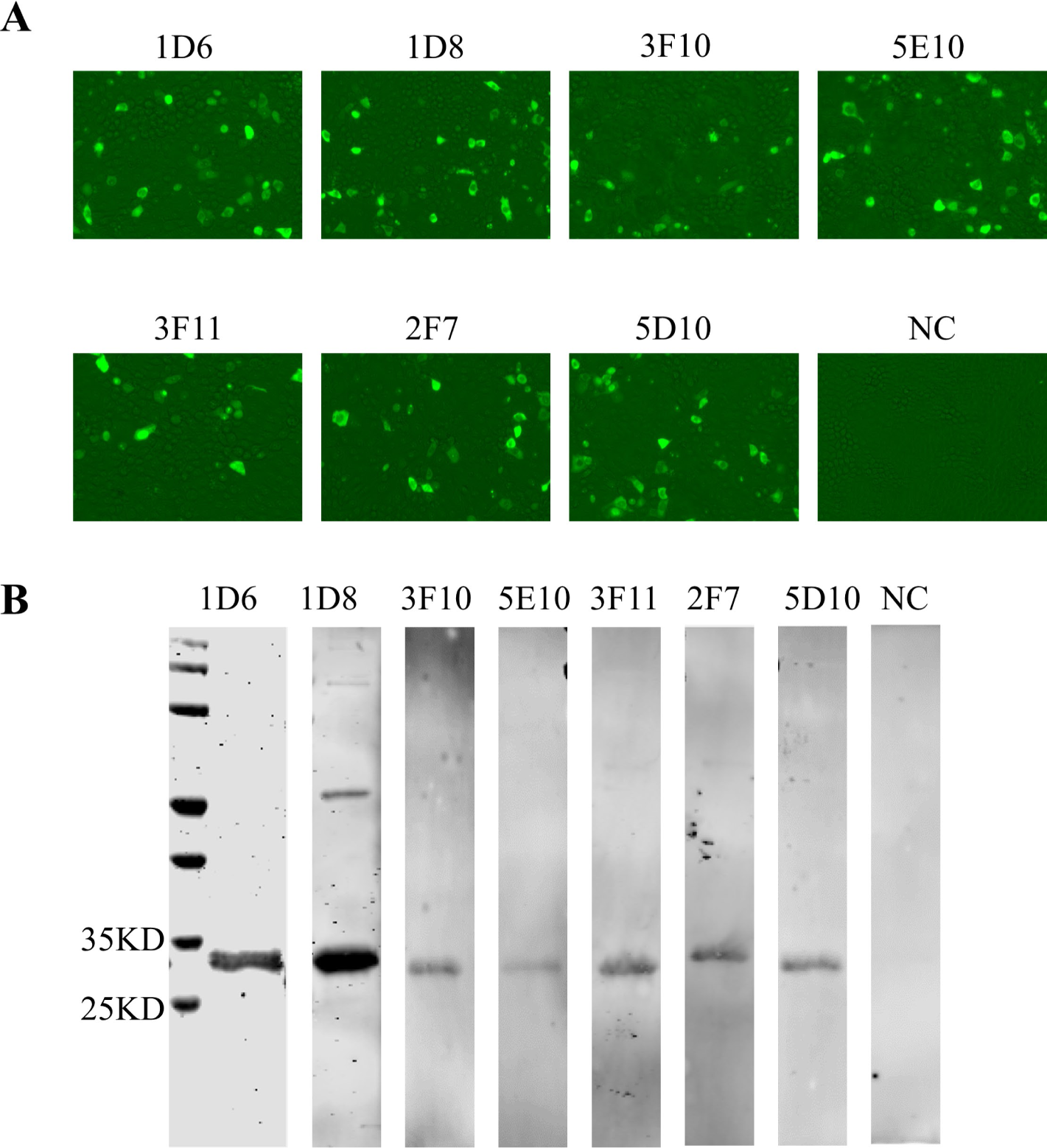
Identification of monoclonal antibodies. (A) Reactivity of PCV4 Cap to MAb 1D6, 1D8, 2F7, 3F10, 3F11, 5D10 and 5E10 in IFA. In 48-well plates, PK-15 cells were transfected with 200 ng of pCI-Cap plasmid in each well. After approximately 24 hours, the cells were fixed and subjected to IFA using monoclonal antibodies and SP2/0 cell supernatant. (B) Characterization of mAbs by Western blot. The purified PCV4 VLPs were subjected to SDS-PAGE and analyzed by Western blot. MAb 1D6, 1D8, 2F7, 3F10, 3F11, 5D10, and 5E10 were used as the primary antibodies, and HRP-conjugated goat anti-mouse IgG (Sigma, diluted 1:10,000, USA) was used as the secondary antibody.

### Identification of the epitope recognized by the MAb 1D8

To map the epitope recognized by seven MAbs on the capsid of PCV4, we expressed GST-fused overlapping peptide fragments using the vector pGEX-6p-1. We then assessed the reactivity of these peptides with MAbs by Western blot. The result revealed that all seven monoclonal antibodies recognize the same epitope (Data not shown). We chose the MAb 1D8 as a representative.

Initially, we expressed three overlapping peptides spanning the full-length Cap protein of PCV4 (Fig.3A-B). The epitope was located within the GST-Cap162-228 based on its reactivity with MAb 1D8 (Fig.3C). To pinpoint the exact peptides binding to the MAb 1D8, we expressed four overlapping peptides spanning the GST-Cap162-228 (Fig.3D-E). The results showed that the MAb 1D8 reacted with the GST-Cap 214-228, suggesting the epitope recognized by the MAb 1D8 located within the 214-228 of the Cap protein (Fig.3F).

**FIG 3.**
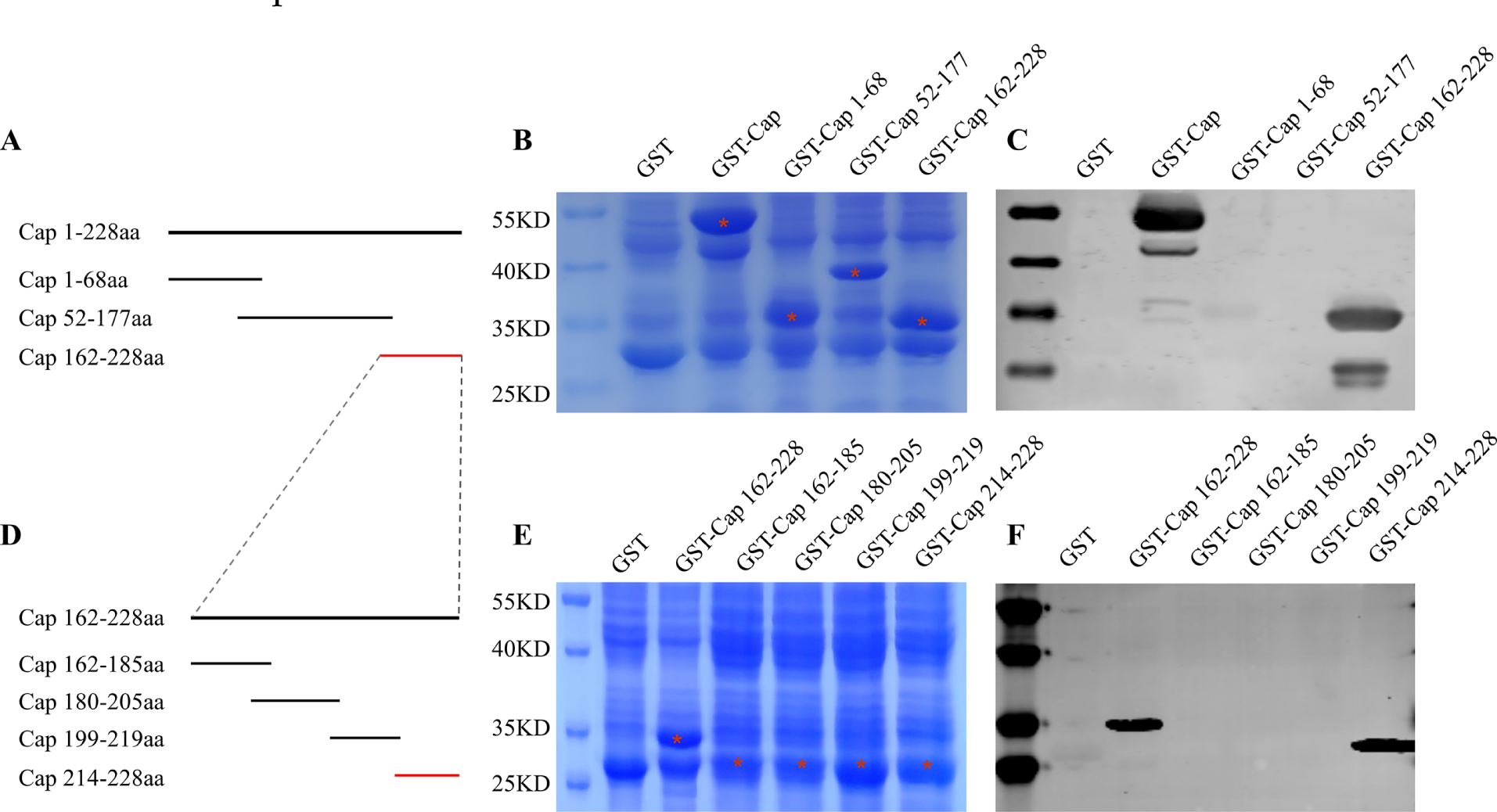
Epitope Mapping of MAb 1D8. (A) Schematic representation illustrating the expression strategy used for three overlapping truncated fragments of the Cap protein (Cap1-68aa, Cap52-177aa, and Cap162-228aa). (B) The three truncated Cap fragments were expressed in E.coli in nearly equimolar amounts, then subjected to SDS-PAGE analysis and indicated with red asterisks. (C) The reactivity of these truncated Cap fragments with MAb 1D8 was assessed by Western blot. GST-Cap was used as a positive control, and GST was used as a negative control. (D) To refine the localization of the epitope within the Cap162-228aa segment, four overlapping truncated fragments of Cap162-185aa, Cap180-205aa, Cap199-219aa, and Cap214-228aa were expressed. (E) The four truncated fragments were expressed in E.coli in approximately equimolar amounts, then subjected to SDS-PAGE analysis and indicated with red asterisks. (F) The reactivity of these truncated Cap fragments with MAb 1D8 was analyzed by Western blot. GST-Cap162-228aa was used as a positive control, and GST was used as a negative control.

To precisely map the minimal epitope recognized by the MAb 1D8, we used a stepwise reduction of individual amino acids from the N-terminus of the GST-Cap 214-228 peptide (Fig.4A-B). GST fusion peptides were expressed (Fig.4C). The MAb 1D8 still reacted with P12-15, which had been progressively truncated from the N-terminus and shortened by 11 residues. However, P10-14, which had only been truncated by one residue from the C-terminus, lost its reactivity (Fig.4D). Therefore, P12-15 (Cap225-228aa) is the minimal epitope recognized by MAb 1D8.

**FIG 4.**
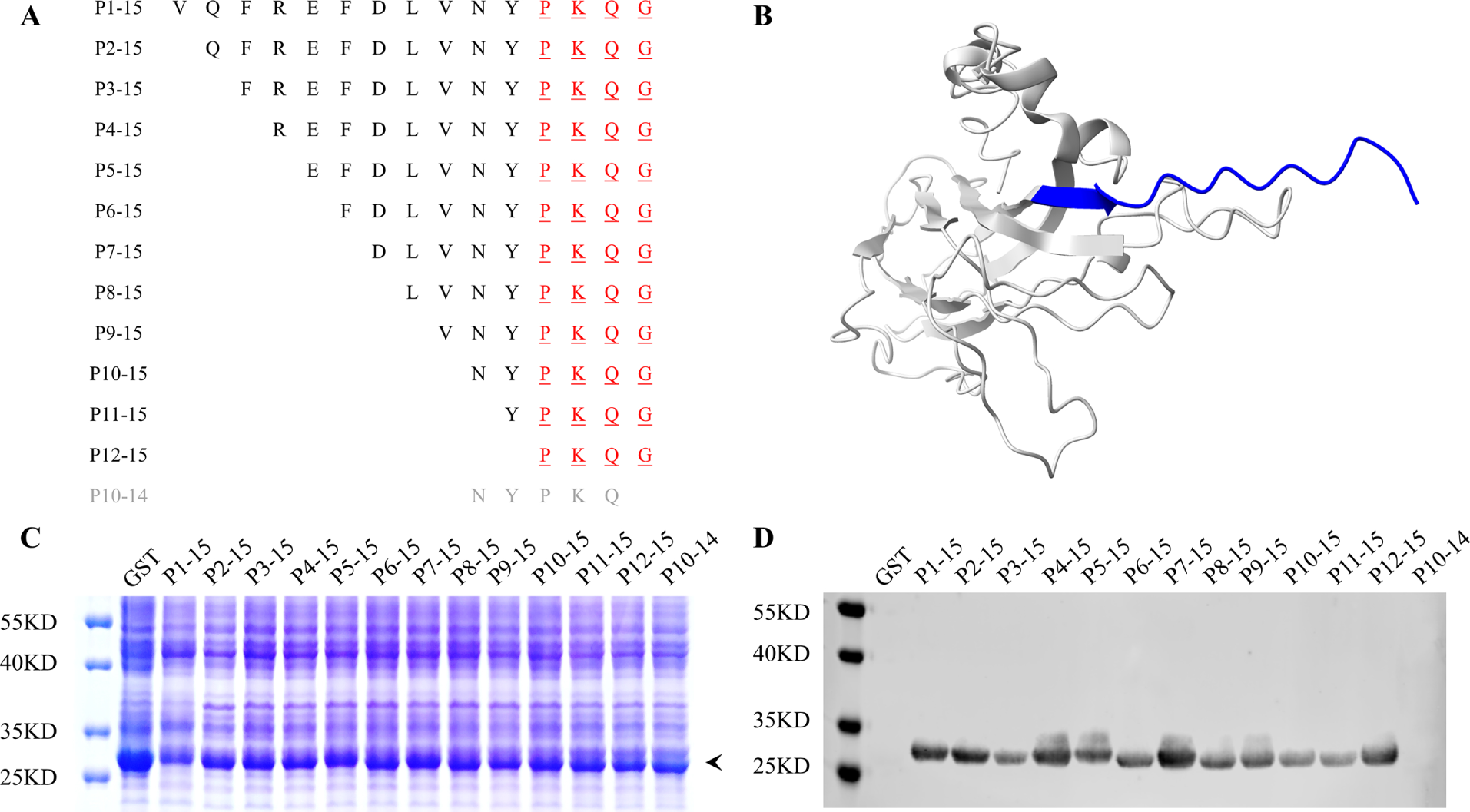
The minimal epitope reacts with MAb 1D8 in the Cap 214-228aa fragment. (A) To identify the minimal epitope recognized by MAb 1D8 within the Cap 214-228aa fragment, we performed successive truncations of individual amino acids from both the N-terminus and C-terminus of the GST-Cap 214-228 peptide. This resulted in the expression of 12 GST fusion peptides. The expression strategy is depicted as shown. (B) The diagram illustrates the location of the Cap 214-228 peptide in Cap protein, highlighted in blue. (C) Nearly equal amounts of 12 truncated GST-Cap fragments were expressed in E. coli. The lysate supernatant was collected and analyzed using SDS-PAGE. The black arrow indicates the truncated GST-Cap peptides. (D) The reactivity of these truncated GST-Cap peptides with MAb 1D8 was analyzed by Western blotting. Peptide P1-15 was used as a positive control and GST as a negative control.

### Critical amino acids recognized by the MAb 1D8

To identify the key residues of the epitope recognized by the MAb 1D8, we replaced each of the nine residues of the core sequence ^220^DLVNYPKQG^228^ with Ala and subjected them to Western blot (Fig.5A-D). Replacing any of the amino acids (Lys^226^, Gln^227^, Gly^228^) in the core sequence completely abolished the binding activity of the peptide to the MAb 1D8, indicating that the motif ^226^KQG^228^ plays a critical and indispensable role.

**FIG 5.**
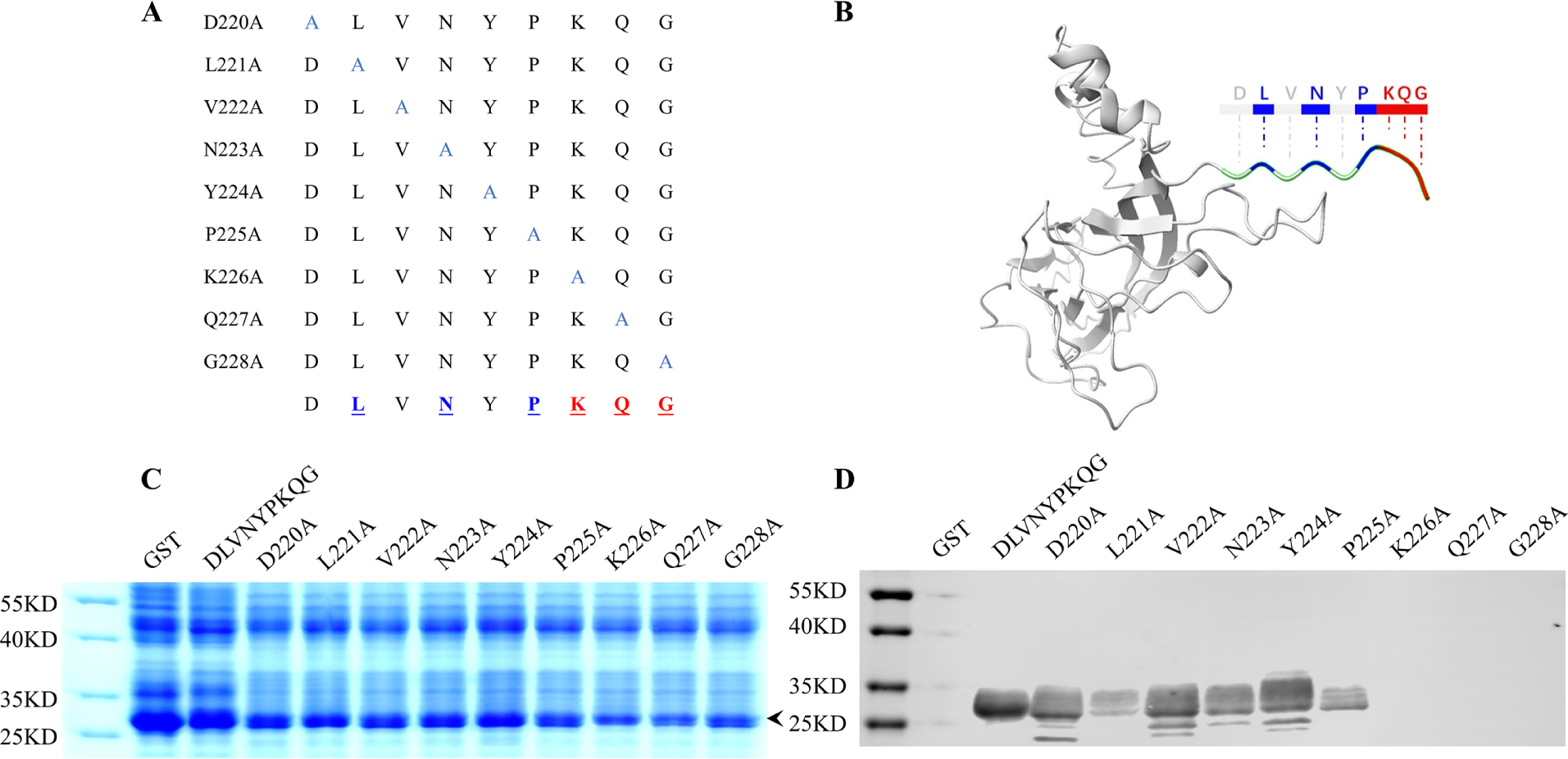
Alanine scanning to determine key amino acids in the ^220^DLVNYPKQG^228^. (A) This schematic illustrates the mutation strategy. (B) The model displays the location of the 220DLVNYPKQG228 epitope and its key amino acids. (C) Nine mutated GST-fusion proteins were expressed in E. coli. Approximately equal amounts of monoalanine-displaced GST-fusion peptides were subjected to 12% SDS-PAGE gels, indicated with a black arrow. (D) The mutant fusion proteins were transferred onto a PVDF membrane and tested for reactivity using MAb 1D8. The unmutated GST-^220^DLVNYPKQG^228^ peptide was the positive control, while GST was the negative control.

Additionally, the binding activity of the peptide to the MAb 1D8 significantly decreased after substituting three amino acids (Leu^221^, Asn^223^, Pro^225^), suggesting that Leu^221^, Asn^223^, and Pro^225^ play crucial roles in the affinity of the epitope to the MAb 1D8.

### Spatial localization of the epitope

The PCV4 Cap protein sequence was used for homology modeling to obtain its monomer protein and the VLP structure. The epitope ^225^PKQG^228^ is located in a random coil within the last four residues of the C-terminus using ChimeraX. It extends outward from the surface of both Cap and VLP (Fig.6A). Moreover, the immunoelectron microscopy result showed that the MAb 1D8 could bind to the surface of VLP and was labeled by a gold-labeled secondary antibody (Fig.6B). These results confirmed that the identified epitope is on the outer surface of Cap and VLP.

**FIG 6.**
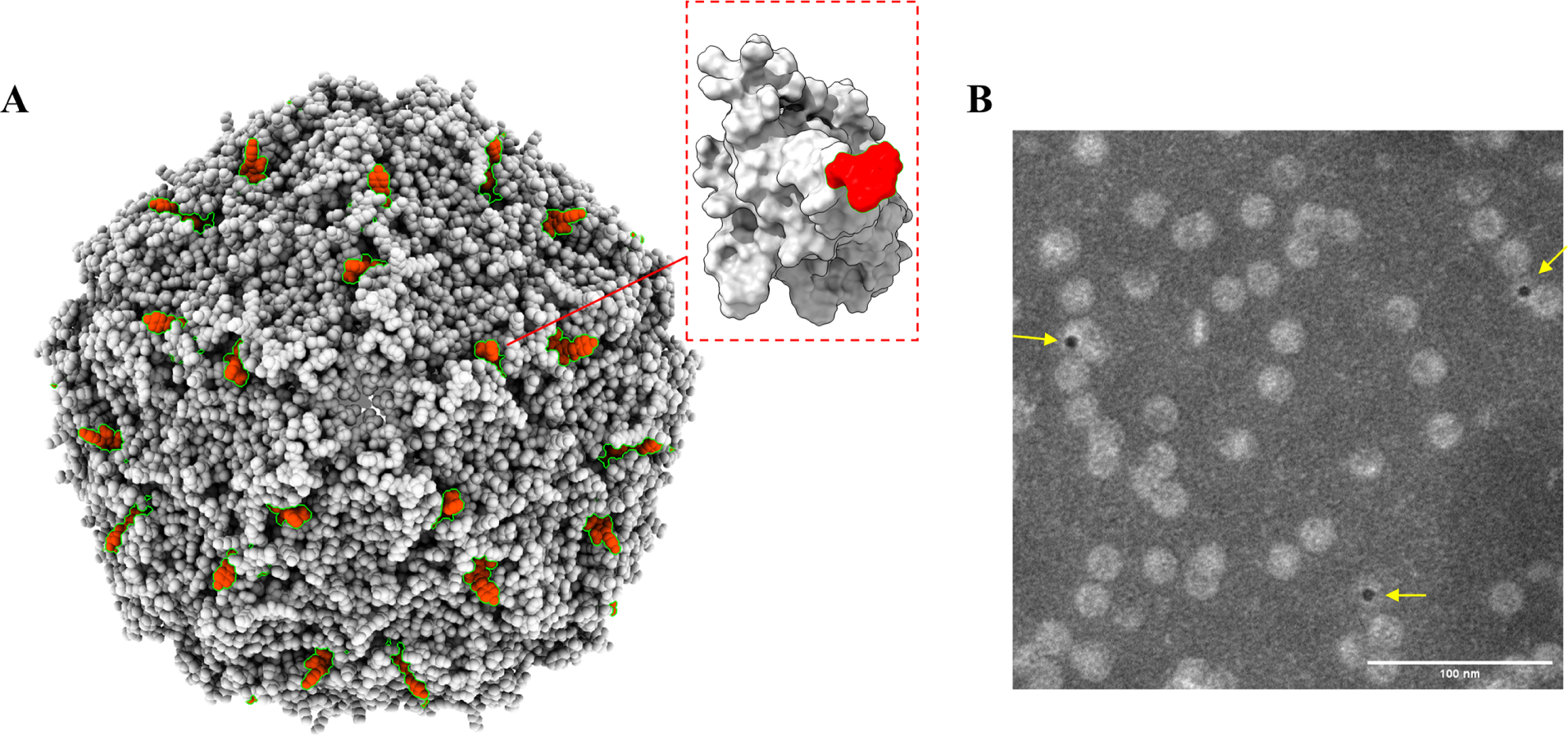
The spatial structural position of the recognition epitope for mAb 1D8 on the Cap protein and VLP. (A) In the homology modeling plot of PCV4 Cap protein and VLP, we observed that the recognized epitope extends out of the outer surface of the VLP. The epitope is marked in red. (B) Immunoelectron microscopy observations were conducted using MAb 1D8 as the primary antibody and 5 nm colloidal gold-conjugated antibody as the secondary antibody for detection. TEM revealed gold labeling on the surface of VLPs.

### PCV4-Specific epitope recognized by the MAb 1D8 is highly conserved

Since the MAb 1D8 reacts specifically with the PCV4 Cap protein, sequence alignment of the epitope with ten PCV4 Cap proteins retrieved from GenBank was performed. The examination revealed that the critical residues ^225^PKQG^228^ recognized by the MAb 1D8 are highly conserved among all ten PCV4 strains, with no observed mutations. Furthermore, the Cap proteins from other PCV genotypes do not contain the MAb 1D8-recognized peptide (Fig.7A). The absence of cross-reactivity with PCV2 and PCV3 counterparts was confirmed by Western blot, which verified the exclusive reactivity of MAb 1D8 with PCV4 Cap proteins (Fig.7B). A BLAST search for the sequence ^225^PKQG^228^ in Genebank yielded no occurrences of the identified peptide in other porcine pathogens except PCV4 Cap. We concluded that the MAb 1D8 can be a valuable tool for accurately diagnosing PCV4 without detecting other PCV genotypes.

**FIG 7.**
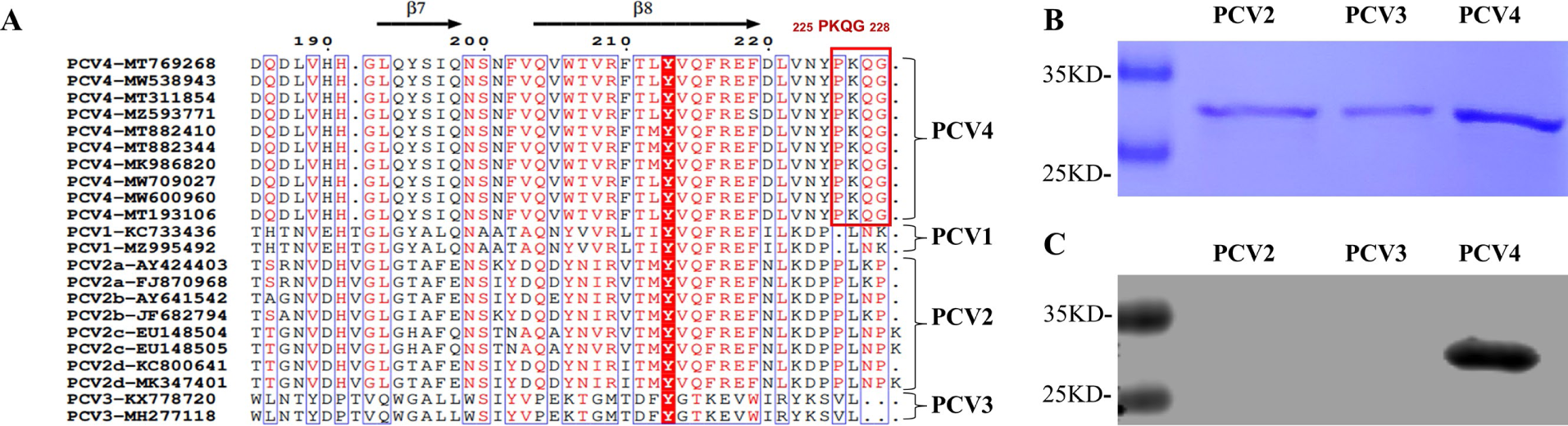
Analysis of the conservation of the ^225^PKQG^228^ epitope in PCV4. (A) The amino acid sequences of 10 PCV4 Cap proteins and 12 other PCV Cap proteins retrieved from GenBank were comparatively analyzed for the identified epitope. (B) The laboratory-stored purified PCV2-4 Cap proteins were subjected to SDS-PAGE. (C) Western blot probed the reactivity of MAb 1D8 with PCVs. MAb 1D8, which was identified as reactive with the ^225^PKQG^228^ epitope, was used as a primary antibody.

## DISCUSSION

PCV4 is an emerging circovirus identified in 2019, and its current information is very limited[6]. For detection and potential future vaccine design, it is important to focus on studying the immunogenicity and epitopes of PCV4 Cap. Extensive studies of PCV2 can serve as references for PCV4 epitopes and immunogenicity studies. Although PCV4 differs in sequence and their surface loop regions from PCV2[12, 30], they share structural similarities. Here, seven MAbs were screened and generated by immunization with VLPs, which closely mimic the structure of the PCV4 original virus[23, 25, 30, 38]. The antigenicity and immunogenicity of PCV4 VLP are similar, closely mirroring the natural immune response to the virus in vivo[39]. Also, we identified a new PCV4 epitope located at the C-terminus of the Cap protein and determined it is on the outer surface of VLP. This is the first analysis and report of PCV4 epitope, which proved to be PCV4-specific and highly conserved. Our results contributed to immunogenicity and future studies on PCV4. The MAb 1D8 can be used to develop diagnostic tools specifically detecting PCV4, which is valuable for future epidemiological distribution studies.

The computational prediction of B-cell epitopes showed eight potential epitopes. Most of them are located on the loop structure. However, relying on a single method for epitope exploration has limitations[40]. We finely mapped an epitope ^225^PKQG^228^ of PCV4 recognized by the MAb 1D8 using the truncated protein expression and alanine scanning mutagenesis method. Seven MAbs were found to bind to the same site, specifically residues 214-228 at the C-terminus (Data not shown). The epitope ^225^PKQG^228^ is the last four residues of the PCV4 Cap protein. This is the first report on PCV4 epitope identification. Based on previous studies on PCV2, the C-terminal peptides play essential roles in the evolution of PCV, pathogenesis, and proliferation[39, 41, 42]. For instance, PCV2 interacts with proteins containing the SH3 structural domain through PXXP, thereby affecting viral replication[42, 43], indicating that the last four C-terminal residues form the highly conserved motif PXXP. This position in PCV2 is not only a linear epitope but also participates in the constitutive conformation[41, 44, 45]. In this study, the identified ^225^PKQG^228^ could also be a linear epitope on PCV4. Moreover, the antibodies generated by epitope at the C-terminus of PCV2 can differentiate between natural infection and vaccine immunization[39]. Regarding previous studies on PCV2, it is hard to ignore the importance of the PCV4 conserved epitope ^225^PKQG^228^, which was identified at the corresponding position in the present study. Further research is required to explore the role of this epitope during PCV4 infection and its application.

It has been demonstrated that epitope ^231^LNP^233^ at the C-terminus of PCV2 serves as a common neutralization epitope for PCV1 and PCV2[41]. As it is located in the corresponding position on PCV4, further exploration is required to determine whether the epitope ^225^PKQG^228^ exhibits neutralizing activity. The residues Lys^226^, Gln^227^, and Gly^228^ were identified as indispensable in ^220^DLVNYPKQG^228^, as they lost reactivity with MAb 1D8 when mutated to Ala. The mutation of Leu^221^, Asn^233^, and Pro^225^ weakened the reactivity. The affinity of MAb 1D8 to the residues in this epitope alternates. This result is consistent with the structural features of the loop, with Leu^221^, Asn^233^, Pro^225^, Lys^226^, Gln^227^, and Gly^228^ located on the exposed side of the loop (Fig.5A).

The spatial location of the PCV2 C-terminus epitope has been determined to extend out and away from the surface of the virus particle[42, 46]. Based on the PCV2 structure, homology modeling of the PCV4 VLP structure also suggests that the ^225^PKQG^228^ is exposed on the surface of PCV4 (Fig.6A). Furthermore, we showed that the MAb 1D8 bound to the surface of PCV4 VLP by immunoelectron microscopy (Fig.6B). The ability to bind to the surface epitope of PCV4 VLP brings a significant advantage for MAb 1D8 as a detection antibody. Based on the available information, PCV4 is classified into only two genotypes: PCV4a and PCV4b[10, 11, 19]. Upon comparing the sequences of representative PCV1-4 strains, it can be concluded that the epitope ^225^PKQG^228^ is highly conserved in PCV4 and is not present in the amino acid sequences of the other three PCVs (Fig.7A). Additionally, we found that the MAb 1D8 reacted specifically binding to PCV4 Cap and did not cross-react with PCV2 and PCV3 by Western blot (Fig.7B). We are convinced that the MAbs prepared in this study have significant value for the specific diagnosis of PCV4.

In summary, we generated MAbs against PCV4 VLPs and identified a highly conserved epitope on the C-terminus of PCV4 Cap protein. The epitope was finely mapped using the MAb 1D8, and its essential residues were also identified. Further studies should be conducted to investigate the potential biological significance, including whether it is a neutralizing epitope and its contributions to the PCV4 assembly and replication.

## ACKNOWLEDGMENTS

We would like to express our gratitude to our colleagues and lab mates for providing us with such technical support and valuable suggestions from time to time.

## CRediT author statement

**Zheng Fang**: Investigation, Writing-Original Draft.

**Mingxia Su**: Formal Analysis, Validation.

**Xuehui Cai**: Resources.

**Tongqing An**: Project administration, Funding acquisition.

**Yabin Tu**: Conceptualization.

**Haiwei Wang**: Methodology, Conceptualization, Writing-Reviewing and Editing, Supervision, Funding acquisition.

